# Effects of Inaccurate Response Function Calibration on Characteristics of the Fiber Orientation Distribution in Diffusion MRI

**DOI:** 10.1101/760546

**Authors:** Fenghua Guo, Chantal M.W. Tax, Alberto De Luca, Max A. Viergever, Anneriet Heemskerk, Alexander Leemans

## Abstract

Diffusion MRI of the brain enables to quantify white matter fiber orientations noninvasively. Several approaches have been proposed to estimate such characteristics from diffusion MRI data with spherical deconvolution being one of the most widely used methods. Constrained spherical deconvolution requires to define – or derive from the data – a response function, which is used to compute the fiber orientation distribution (FOD). This definition or derivation is not unequivocal and can thus result in different characteristics of the response function which are expected to affect the FOD computation and the subsequent fiber tracking. In this work, we explored the effects of inaccuracies in the shape and scaling factors of the response function on the FOD characteristics. With simulations, we show that underestimation of the shape factor in the response functions has a larger effect on the FOD peaks than overestimation of the shape factor, whereas the latter will cause more spurious peaks. Moreover, crossing fiber populations with a smaller separation angle were more sensitive to the response function inaccuracy than fiber populations with more orthogonal separation angles. Furthermore, the FOD characteristics show deviations as a result of modified shape and scaling factors of the response function. Results with the in vivo data demonstrate that the deviations of the FODs and spurious peaks can further deviate the termination of propagation in fiber tracking. This work highlights the importance of proper definition of the response function and how specific calibration factors can affect the FOD and fiber tractography results.

## 1. Introduction

Diffusion MRI allows to characterize tissue microstructure in vivo and noninvasively by measuring the anisotropic diffusion of water molecules [1, 2]. Diffusion tensor imaging (DTI) [3] is the most widely used model in clinical studies to relate the diffusion MRI signals to the diffusion characteristics of the underlying tissue. However, DTI is inadequate to estimate the directional information in voxels containing crossing fibers [4, 5]. A commonly used approach to resolve more complex fiber configurations in the brain is spherical deconvolution (SD) [6–8]. SD also allows for the extraction of fiber population specific microstructural measures derived from the magnitudes of the fiber orientation distribution (FOD) functions, such as apparent fiber density (AFD) [9] and hindrance modulated orientational anisotropy (HMOA) [10].

SD requires an appropriate response function as input to estimate the FOD [7]. The response function, representing the diffusion signal for a single fiber population, is ideally calibrated from the acquired diffusion MRI data [11, 12]. In brief, for each subject, the voxels containing only single fiber populations are localized, and an average of the diffusivity characteristics within those voxels is used to represent the subject specific response function. An inadequately chosen response function can affect the quantification of FOD characteristics like AFD and HMOA, as well as the fiber tractography.

In order to compare inter-subject AFD, Raffelt and colleagues [9] chose a response function common to all subjects to minimize the differences between subjects for voxel-wise AFD comparison. However, this may potentially result in a bias in the estimated FOD. Specifically, the use of such a common response function for group-wise analysis may cause biases in the FOD peak orientations for individual subjects. Therefore, whereas a common response function is optimal for the comparison of AFD and HMOA in group studies [9], it is unclear whether this is also optimal for group-wise tractography studies because of the potentially inaccurate FOD peak orientations and concomitant spurious FOD peaks. Intuitively, the difference in response function characteristics across healthy subjects are not expected to be large, as response functions are generally averaged from more than hundreds of voxels that are supposed to contain single fiber populations [6,7,12]. This was partly demonstrated by Jeurissen and colleagues [13], who studied the inter-subject response functions of 100 healthy subjects from the Human Connectome Project (HCP) [14] and observed only subtle differences. Accordingly, it seems justified not to be too concerned about inter-subject response function variability in healthy subjects, since either using averaged response functions or individual response functions is not likely to affect the FOD profiles in the HCP dataset. However, although the differences in the response functions of healthy subjects may be small [13], this is likely not the case for subjects with some form of pathology. The inter-subject signal deviations do raise concern for aging and diseased groups.

White matter degeneration, whether caused by aging or by a disease process, may substantially alter the response function. Hence, studying subjects of different ages with a common response function might introduce errors due to discrepancies in white matter characteristics. Therefore as the focus of this work, it is useful to investigate such differences in response functions and the resulting variations of the FOD. A thorough numerical evaluation focusing on the angular characteristics of FOD is needed to shed more light on this issue.

Previous studies have discussed the effect of improperly calibrated response functions on the FOD characteristics and fiber tracking. Tournier [7] and Dell’Acqua [8] reasoned from a mathematical point of view that a wrongly chosen response function would affect the magnitudes of FOD peaks, thus also AFD and HMOA, but would leave their orientations unaffected. Dell’Acqua and colleagues [8, 10] investigated with simulations and in vivo data the effects of various response function changes on the FOD profiles, including variations in the response function shape and scaling factor, as well as in axonal radius and in angle of crossing pathways for the damped Richardson-Lucy (dRL) method. Their paper focused on the effect of the response function on FOD amplitudes and the sensitivity of HMOA to diffusivity changes per fiber population, as compared to traditional metrics as fractional anisotropy (FA) and mean diffusivity (MD). Parker [15] studied the FOD peak orientations and the existence of spurious peaks in simulations as a function of the response function miscalibration for CSD and dRL. The results of that study demonstrate that sharper response functions resulted in more spurious peaks in the FOD profiles, and that the mismatch of the calibrated-targeted response functions introduced uncertainty on the main FOD peak orientations. However, in previous work[15], the authors used the FA value as a metric to characterize the response functions, a strategy which is unable to describe the true axial and radial diffusivities in crossing fibers [16]. Changes in FA entangle changes in the axial and radial diffusivities, so that the effects on these two diffusivities could not be studied straightforwardly. Here we seek to disentangle these effects and, complementing earlier studies [15, 17], also aim to quantify both the effect on peak magnitude and angular deviation.

In this manuscript we studied how variations in the response function affect voxel-wise FOD characteristics and fiber tracking. Changes in pathology are likely reflected in changes in either the axial or the radial diffusivity, which in our study, is represented by the shape and scale factor of the response function. Simulations were designed to explore the effects of the response function shape and scaling factor on the FOD properties, such as the number of FOD peaks, their orientation (for tractography) and magnitude, and the AFD. Additionally, in vivo data from the Human Connectome Project (HCP) were used to illustrate how the choice of the response function in CSD can affect the FOD quantification and fiber tracking.

## 2. Methods

In Sections 2.1 and 2.2, we give a brief background on (constrained) spherical deconvolution methods to reconstruct the FOD. In Section 2.3 we outline the simulation experiments and introduce the shape and scaling factor that characterize the response function. Section 2.4 presents the parameter settings used in these simulations. In Section 2.5, the in vivo data experiments are described.

### 2.1 Constrained Spherical Deconvolution

Recent studies showed that crossing fibers account for over 90% of white matter voxels [4]. The DTI representation cannot resolve crossing fibers by design and thus provides non-specific metrics in such voxels. Spherical deconvolution approaches [6–8,18,19] overcome this limitation and allow for estimating the FOD for more complex fiber configurations, while retaining reasonable computation and acquisition time compared with other methods [20–23].

CSD assumes that the diffusion MRI signals can be expressed as the spherical convolution of a fiber response function with the FODs in the spherical harmonics basis, thus also assuming the validity of the response function in all voxels. The response function represents the diffusion-weighted signal of a single fiber population. Spherical harmonics form a complete basis on the sphere. However, to fully reconstruct a signal on the sphere, the spherical harmonics should have infinite order, which is not possible in practice. In clinical studies, signals with up to 60 gradient directions are generally acquired, limiting the order of the spherical harmonics to 8, which we also adopted in this work.

The FODs are used to infer information on the orientation of the fiber pathways under the assumption that the FOD peak orientations coincide with the underlying fiber directions. To reconstruct the FOD, truncation of the spherical harmonics is needed, causing the so-called “ringing” effect on the FOD profiles, which introduces implausible negative values. In order to suppress the ringing effect and the sensitivity to noise, the regularization of FOD was proposed [7,19,24] to improve the conditioning of the deconvolution problem, which is further referred to as constrained SD (i.e., CSD). In addition to directional information, the magnitudes of the FOD are used to compute additional metrics, such as AFD [9] and HMOA [10]. The accurate estimation of FOD peak directions and magnitudes is therefore essential for subsequent analysis.

### 2.2 Shape and scaling of response functions

The response function used in the CSD process can be either simulated or derived directly from the data. Following the latter approach, which is more common, voxels that have a high chance of containing single fiber populations are used to calibrate the response function. A straightforward approach to numerically implement the concept of a single fiber population is to threshold, for instance, the fractional anisotropy (FA), above a pre-defined value. However, the choice of FA threshold is not trivial and can cause inaccuracies in the response function estimation [12]. A data-driven method using a recursive calibration framework was proposed to estimate the response function from the subject data in an unbiased way [12]. This method estimates which voxels contain single fiber populations by iteratively excluding voxels which do not have a single dominant orientation and updating the estimated response function.

The choice of the fiber response function has an impact on the peak directions and magnitudes of the FODs [10,15,19]. Theoretically, changes in the response function are directly reflected in the FOD estimation, but should affect only peak magnitudes while leaving their orientations untouched [6, 10]. However, in practice, due to the low SNR level in diffusion-weighted MRI data, the ill-posedness of inverse problems, and the regularization process, the effects of the choice of response function on the FODs become less obvious.

Parker et al. [15] investigated alterations of response function by changing its FA value. Here, we acknowledge that changing the FA affects both the scale and the shape of the response function. It is thus not straightforward to disentangle an FOD change into scale and shape effects. To this end, we decompose general changes in the response function into specific changes in shape and scale [8] and analyze their individual effects on the FOD characteristics (i.e., magnitude, the number of peaks, and peak orientations). The following sections describe how we can achieve such changes in shape and scale of the response functions in the simulated and in vivo diffusion MRI data experiments.

### 2.3 Simulation experiments

#### 2.3.1 Modeling of single fiber populations and response functions

If the diffusivity *D* associated with the underlying fiber population is expressed by an axially symmetric diffusion tensor, whose first eigenvector is in parallel with the *z*-axis in the reference coordinate frame, then *D*(*θ, φ*) can be written as (Anderson 2005)

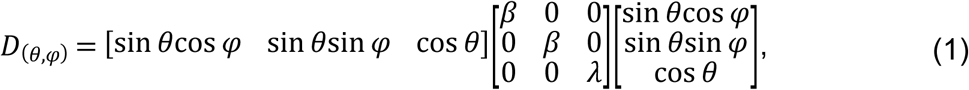

where λ and *β* are the axial and the radial diffusivity of the single fiber population, (*θ, φ*) is the polar angle set between the fiber orientation and the applied gradient. Given the axial symmetry property of the diffusion tensor, Eq. (1) can be simplified as

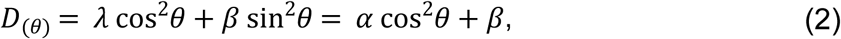

where α = λ - β is the absolute difference between the axial and radial diffusivity. For simplicity, if we assume that the signal *S*(*θ, φ*) from each fiber population is a function of *D*(*θ, φ*), then the diffusion-weighted signal *S* can then be rewritten as [3]

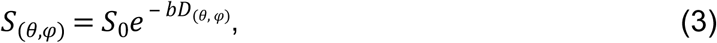

where *S*_0_ is the non-diffusion-weighted signal and *b* is the b-value that represents the strength of diffusion weighting. Combining Eq. (1) – Eq. (3), the diffusion-weighted signals can be expressed as [18]

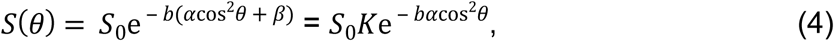

Where *K* = *e^-bβ^* Eq. (4) highlights the dependency of *S* on the shape factor *α* and the scaling factor *K*, following the definition in previous studies [8]. In this equation, the scaling factor *K* depends only on the radial diffusivity of the fiber response, representing the isotropic diffusion within the fiber population, whereas the shape factor *α* depends on the difference between the axial and radial diffusivities, representing the anisotropic diffusion within the fiber population.

#### 2.3.2 Modifying the shape and scaling factor of the response functions

Since the response function *R* is intrinsically based on the shape and scaling of the fiber population diffusivities, *R* can be written in the same form as the signal of a fiber population imposed by the gradient at an elevation angle *θ* with the fiber orientation, which is identical to Eq. (4), i.e.,

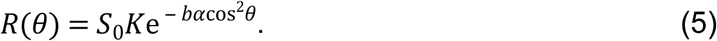

According to Eq. (5), we can modify (i) the shape factor *α* of the response function, by varying only the axial diffusivity with a fixed radial diffusivity, to keep *K* constant; and (ii) the scaling factor *K* of the response function, by changing simultaneously the axial and radial diffusivity, to not alter the shape factor *α*. We can then study the effects of *R* on FOD characteristics, by selectively introducing a discrepancy into the shape or the scale of a simulated single fiber signal with respect to the response function.

#### 2.3.3 Modeling of multi-fiber populations

We model the diffusion-weighted signal within a voxel as the sum of multiple compartments measured from each fiber population. Each compartment is assumed to share an identical response function, so the diffusion-weighted signals are depending only on the orientations of the fiber populations in the voxel and on data noise. We further assume that there is no exchange of water molecules between fiber populations, and that each single fiber population can be represented by a signal *Si*_(*θ*)_ (where *i* denotes the *i^th^* fiber population). The signal *S_DW_* generated by a crossing fiber configuration can then be described by

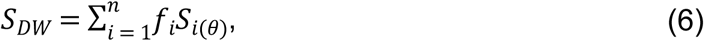

where *f_i_* is the volume fraction of each fiber population, *n* is the total number of fiber populations intercrossing the voxel, and *i*(*θ*) is the angle between the applied gradient and the *i^th^* fiber population. In our work, we focus on configurations of two crossing fiber populations, but the equations of generating the diffusion-weighted signals can also be extended to analyze the FOD characteristics for more than two fiber populations.

#### 2.3.4 Data analysis

Amongst the SD frameworks, the CSD approach is implemented in several software packages, such as *MRtrix* [25], *Dipy* [26] and *ExploreDTI* [27]. In this work, the FODs were estimated with CSD as implemented in *ExploreDTI*. The FOD peak orientations, which are assumed to reflect the underlying fiber orientations [6], and the magnitudes of the FOD peaks, were extracted using a Newton-Raphson gradient descent method [28]. All FOD peaks that were smaller than an absolute threshold of 0.1 were regarded as contributions from noise and thus discarded to reduce false positives [29]. All peaks were clustered to the nearest simulated peak directions, by using an angular threshold of 45° to determine whether or not two peaks were belonging to the same fiber population. In case of simulating multiple fiber populations, only the estimated FOD peaks closest to the simulated fiber populations were considered. For each simulation, the mean and standard deviation of the following FOD metrics were evaluated:

a. the average difference between the estimated and simulated number of FOD peaks;
b. the angular deviations between the estimated FOD peak orientation and the simulated fiber orientation;
c. the estimated separation angles in case of multiple fiber populations;
d. the FOD peak magnitudes in case of single fiber populations;
e. the percentage difference of the estimated AFD with respect to the AFD with the reference response function.

The AFD computation was performed as the integral of the FOD magnitudes assigned to each peak, which in the literature is commonly referred to as “lobe”. The calculation of the AFD is similar to what was used in a previous study [30], except that we use the gradients generated by the electromagnetic model [31] to segment the FODs for each fiber population instead of using gradients generated by an icosahedron model.

### 2.4 Parameter settings

We simulated different fiber configurations with a predefined *b*-value equal to 3000 s/ mm^2^, a set of 60 gradient directions [31], and *S*_0_ = 1. Rician noise (1000 noise instances) was added to the diffusion weighted signals to simulate SNR (with respect to *S*_0_) levels of [50 40 30 20 15 10]. In the first simulation, a single-fiber configuration was generated with the main diffusion direction along the *z*-axis, setting *α* = 1.2 × 10 ^‒ 3^ mm^2^/s and *K* = 0.4 (i.e. *β* ∼ (0.3 × 10 ^‒ 3^ mm^2^/s)). In the second simulation, a second fiber population was rotated around the *y*-axis and combined with the single-fiber population generated in the first simulation to achieve a separation angle *ω*. Here we simulated crossing fiber populations with separation angles *ω* = [90°, 75°, 60°, 55°, 50°, 45°, 40°].

For both simulations, two sets of response functions were tested to achieve (a) different shape but the same scaling factors, by increasing *α* from 0.6 × 10 ^‒ 3^ mm^2^/s to 1.8 × 10 ^‒ 3^ mm^2^/s with steps of 0.1 × 10 ^‒ 3^ mm^2^/s, while keeping *K* constant (Fig. 1a); and (b) the same shape but different scaling factors, by decreasing *K* from 0.7 to 0.3 with steps of 0.1, while keeping *α* constant (Fig. 1b).

**Fig. 1.**
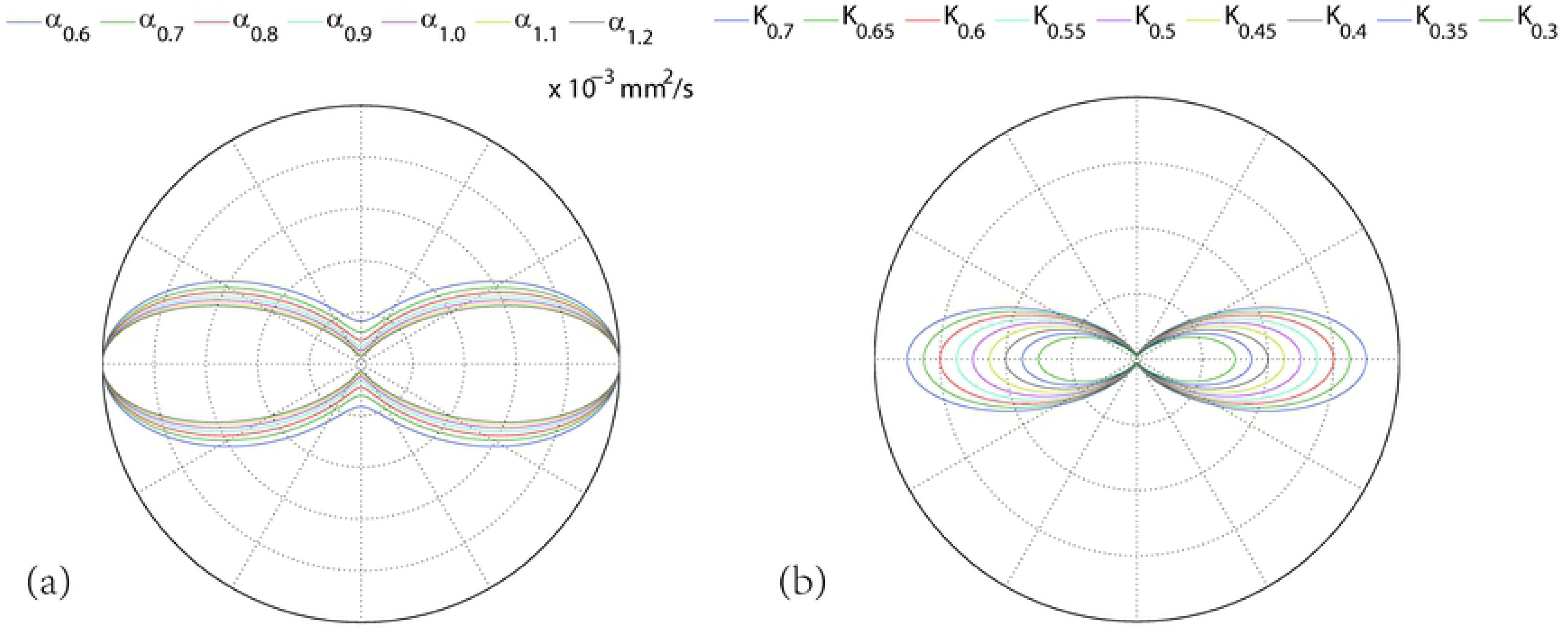
The 2D projection of response functions obtained by changing (a) the shape factor α and (b) the scaling factor *K*. The shape factors are defined from 0.6 × 10 ^‒ 3^ mm^2^/s to 1.8 × 10 ^‒ 3^ mm^2^/s in steps of 0.1 × 10 ^‒ 3^ mm^2^/s. The scaling factors are varied from 0.7 to 0.3 in steps of 0.05.

### 2.5 Peak clustering and angular threshold

We clustered the peak directions to make sure that we are always comparing the angular deviations between the simulated fiber orientation and the FOD peak orientation most closely aligned to that orientation. Like in other studies [16,32,33] that compare axial and radial diffusion characteristics, we also included an angular threshold (e.g., cos (*θ*) > 0.7, which means approximately *θ* < 45°) to make sure the correct peaks were being extracted for further evaluations.

### 2.6 In vivo data experiments

Diffusion-weighted MRI data of a single HCP subject was further used to illustrate the effects of ill-defined response functions on voxel-wise FOD characteristics and brain tractography. In summary, diffusion-weighted images were acquired along 90 diffusion gradient directions with a *b*-value of 3000 s/mm^2^ in addition to 18 non-diffusion-weighted images, and with an isotropic spatial resolution of 1.25 × 1.25 × 1.25 mm^3^. We performed CSD based tractography in *ExploreDTI* with a step size of 1 mm, an FOD threshold of 0.1, an angular threshold of 30°, and seeding points per 2mm × 2mm × 2mm across the whole brain. All the tracts were constructed with deterministic fiber tracking to facilitate data interpretation.

#### 2.6.1 Modeling the response function

The reference response function for the in vivo dataset was represented by the diffusion tensor fit to the response function, as estimated with the recursive calibration approach [12]. Similar to the method described in Section 2.3.2, the diffusion tensor was used to model the changes in the shape and the scaling factor of the response functions. The shape factor *α* of the response function was modified by +/-[0.1 ‒ 0.3 × 10 ^‒ 3^ mm^2^/s], while the scaling factor *K* was modified by +/-[0.1 ‒ 0.2].

#### 2.6.2 Evaluation of in-vivo data

In analogy with the simulations, we computed the voxel-wise difference in number of estimated FOD peaks, the angular deviations of the main orientation, and the percentage difference in AFD of the dominant fiber orientation, for all the estimated FODs. The comparisons of number of FOD peaks were computed for the whole brain, whereas the comparisons of angular deviation and AFD were only computed for voxels with FA > 0.2.

Individual white matter fiber bundles were extracted by using the regions of interest (ROIs) as suggested by Wakana [34]. The segmented fiber pathways include parts of the splenium of corpus callosum (sCC), the genu of corpus callosum (gCC), the cingulum (Cg), the uncinate fasciculus (UF), the corticospinal tract (CST), and the temporal part of the superior longitudinal fasciculus (tSLF). The average FOD characteristics for each fiber bundle were calculated. In addition, FOD characteristics of the response function were computed from (1) the region with a single fiber population as identified during the recursive calibration step (referred to as “SFP-mask”); and (2) the region with voxels for which FA > 0.2 (referred to as “FA-mask”).

## 3. Results

### 3.1 FOD characteristics of single fiber populations

Fig. 2 shows the effect of changing the shape factor and the scaling factor of the response function on the FOD characteristics in a single fiber population. At SNR < 20, the average number of spurious peaks increases when the shape factor increases, but only slightly increases when the scaling factor decreases (Fig. 2A). The angular deviation depends mainly on the SNR and is far less affected by changes in shape or scale factor of the response function (Fig. 2B). By contrast, changes in peak magnitude (Fig. 2C) and the AFD (Fig. 2D) as a function of shape and scaling factor of the response function are more pronounced than due to differences in SNR level alone. Notice that the effect of changing the scaling factor (up to ∼60%) is roughly three times larger compared to changing the shape factor (up to ∼20%).

**Fig. 2.**
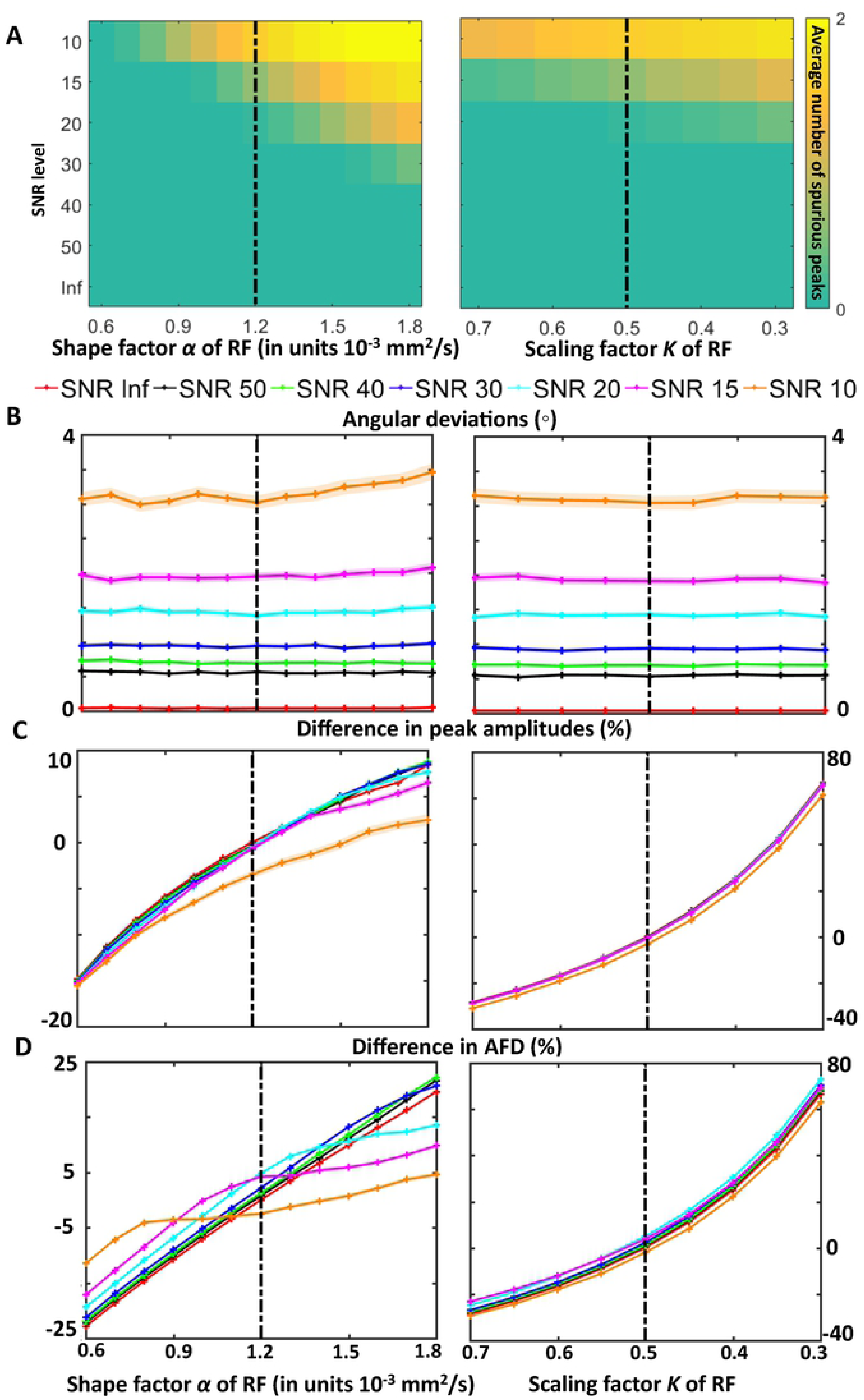
Effect of simulating changes in the response function on FOD characteristics for a single fiber configuration at different SNR levels. Shape factor *α* and the scaling factor *K* of the response function (RF) are varied at different SNR levels to investigate (A) the introduction of spurious peaks, i.e., the average difference between the estimated and predefined number of FOD peaks; (B) the confidence interval (average ± standard error) of the angular deviation of the primary FOD peak; (C) the percentage difference between the amplitudes of the estimated FOD peak and the ground-truth FOD peak; and (D) the percentage difference between the estimated AFD of the primary fiber population and the ground-truth AFD. The dashed vertical lines represent the ground-truth values.

### 3.2 Occurrence of spurious peaks

Fig. 3 shows the average difference between the number of estimated and simulated FOD peaks in relation to the shape (left) and the scaling (right) factor of the response functions for different SNR levels. Overall, performing spherical deconvolution with sharper response functions (i.e., higher values of the shape factor) generally introduces more spurious peaks. On the other hand, CSD fails to extract all the simulated peaks from the estimated FODs when the response function shape factor has smaller values, in particular for separation angles below 55°. With higher noise levels, more spurious peaks are introduced, especially for higher values of the shape factor. Furthermore, adjusting the scaling factor has no significant effect on the estimated number of spurious peaks. While there are hardly any spurious peaks introduced at the lower noise levels (SNR = 30 and 50), additional incorrect peaks can be observed at the higher noise level (SNR = 10).

**Fig. 3.**
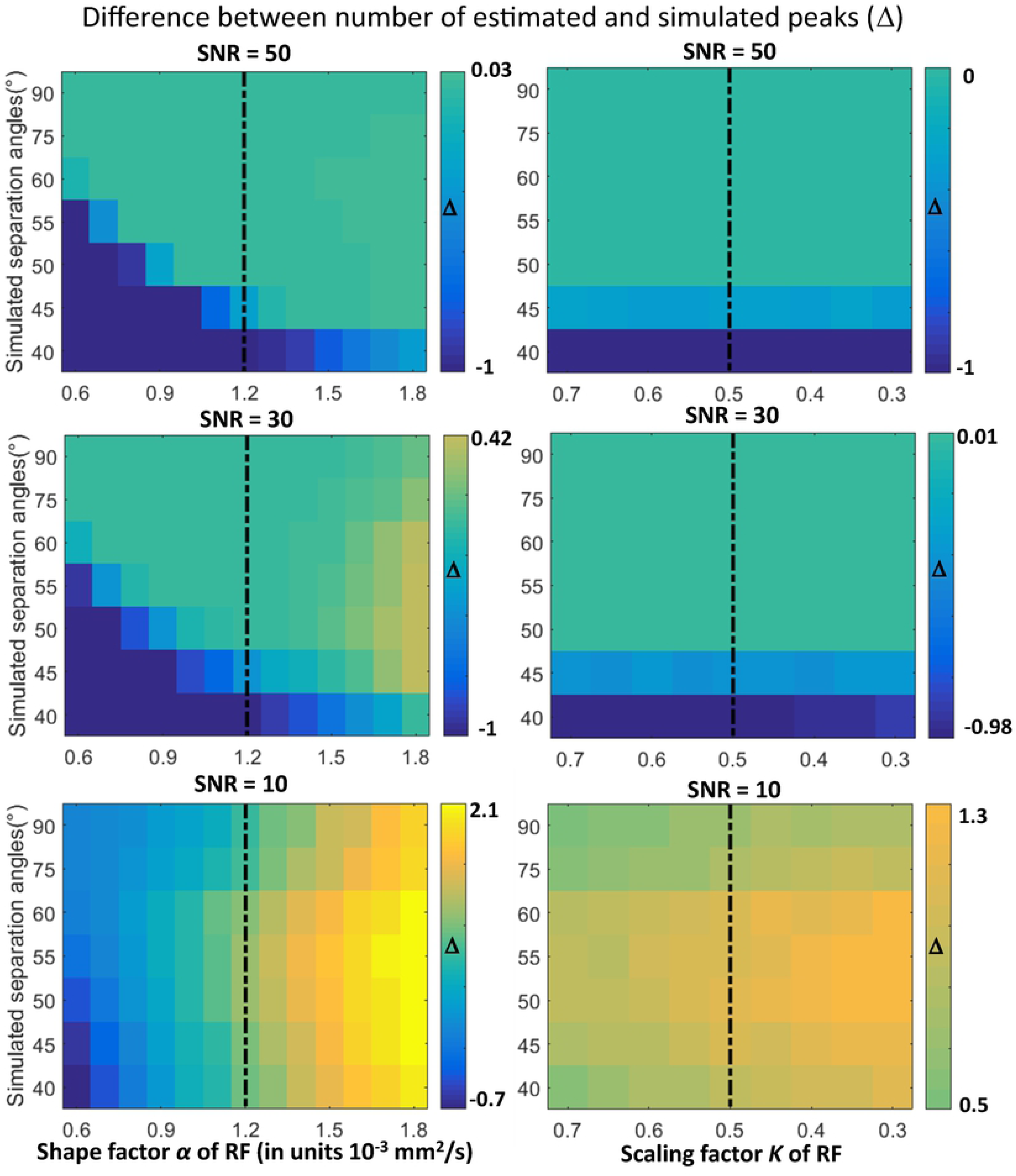
The average difference between the number of estimated and simulated FOD peaks as a function of shape (left) and scaling (right) factor of the response function (RF) at three SNR levels (different SNR for each row). Brighter yellow areas show a higher probability of introducing spurious peaks, whereas darker blue areas show a higher probability of merging the two simulated peaks into one peak. The dashed vertical lines indicate that the settings of the response function are identical to those used for generating the underlying signals. Notice that different scaling of the colorbars were used for better contrast.

### 3.3 Angular deviation

#### 3.3.1 The effect of the shape factor

Fig. 4 shows the results of investigating the effect of the response function’s shape factor on the angular characteristics of FOD peaks at SNR = 50, 30 and 10 for crossing fiber configurations with different separation angles. At lower noise levels (SNR = 30 and 50), lower values of the shape factor generally cause an underestimation of the separation angles, except when the two simulated fiber populations are orthogonal to each other (i.e., 90°) (Fig. 4A). At the higher noise level (i.e., SNR = 10), the bias in the estimated separation angle due to changes in the shape factor is swamped by the noise itself, especially for lower separation angles. From the observed angular deviations in Fig. 4B (the first peak) and Fig. 4C (the second peak) we can observe, in general, that for smaller simulated separation angles, the adverse effects of changing the shape factor of the response function on the estimated FOD angular characteristics are more pronounced.

**Fig. 4.**
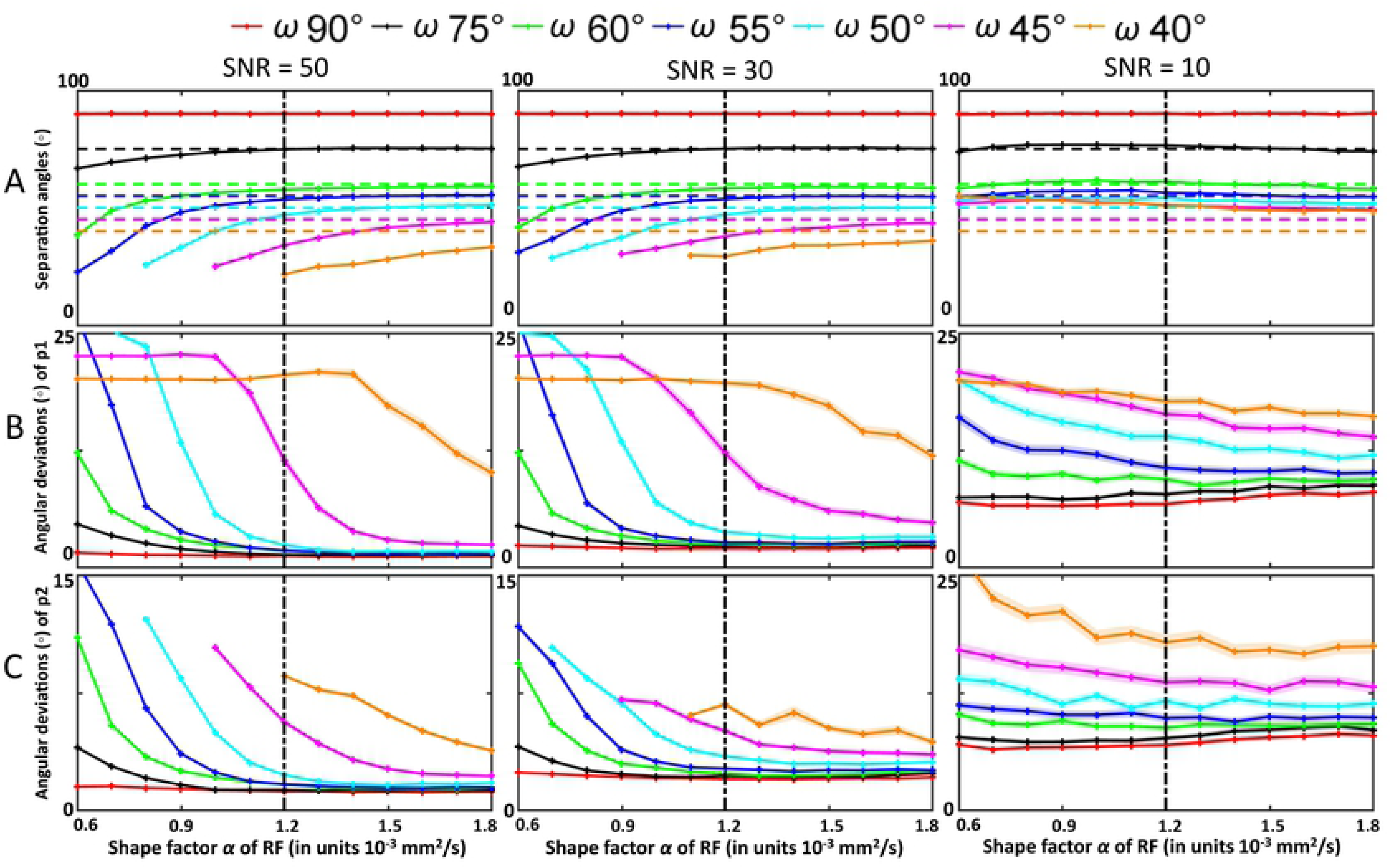
Results of exploring the impact of response functions with different shape factor α on the FOD peaks for crossing fiber configurations simulated with separation angles ranging from 90° to 40°. Fig. 4A shows the estimated separation angles between the two primary peaks. Dashed horizontal lines indicate the simulated separation angles. Fig. 4B and Fig. 4C show the angular deviations between the estimated first (p1) and second (p2) FOD peaks and their corresponding simulated fiber orientations. Solid line interruptions occurred when one of the two peaks was not detected. The means of the estimated values are plotted with the standard error as the shaded areas. Dashed vertical lines are defined as in Fig. 3.

#### 3.3.2 The effect of the scaling factor

Fig. 5 shows the angular deviations between the orientation of the estimated FOD peaks and the simulated fiber orientations as a function of the scaling factor. Overall, crossing fibers with separation angles smaller than 45° show larger angular deviations than those with more orthogonal separation angles. In Fig. 5A, the estimated separation angles do not change significantly as a function of the scaling factor of the response function. Nevertheless, smaller simulated separation angles result in a larger bias of the estimated separation angles. Fig. 5B and Fig. 5C present the angular deviations of the first and second FOD peak, respectively. The angular deviations are not significantly affected by the scaling factor, but do depend on the magnitude of the separation angles of the two fiber populations.

**Fig. 5.**
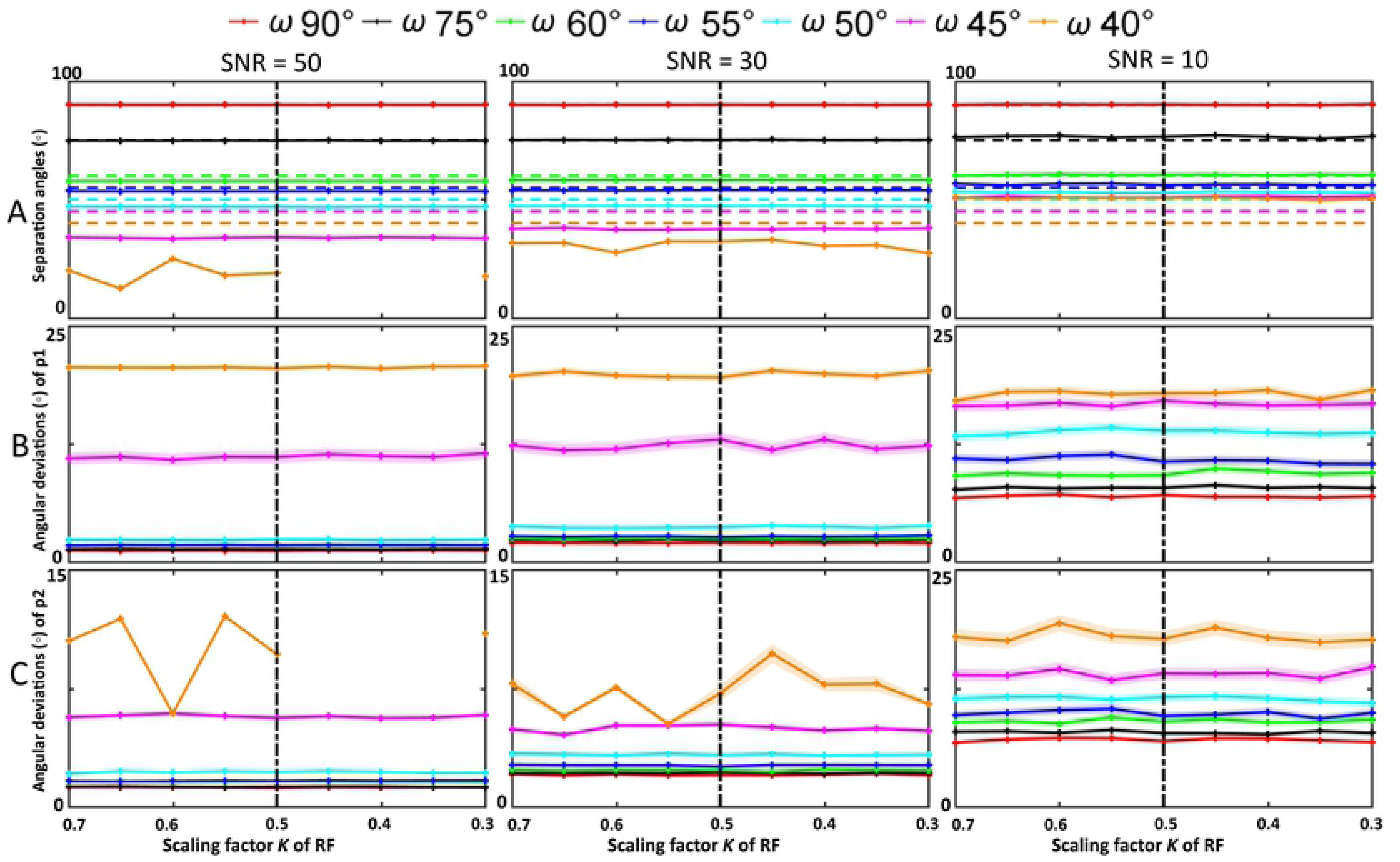
The effect of varying the scaling factor (*K*) of the response function on the FOD peaks for crossing fiber configurations simulated with separation angles ranging from 90° to 40°. Fig. 5A shows the estimated separation angles between the two primary peaks. Dashed horizontal lines indicate the simulated separation angles. Fig. 5B and Fig. 5C show the angular deviations between the estimated first (p1) and second (p2) FOD peaks and the corresponding simulated fiber orientations. Solid line interruptions occurred when one of the two peaks was not detected. The means of the estimated values are plotted with the standard error as the shaded areas. Dashed vertical lines are defined as in Fig. 3.

#### 3.4 AFD per fixel

Fig. 6 shows the percentage difference of the AFD of the first and second fiber population in relation to the response function shape factor (A, B) and scaling factor (C, D). In Fig. 6A, at SNR 50 and 30, the AFD started at a very high value when the shape factor is smaller than 0.8, 1.0 and 1.4 × 10^-3^ mm^2^/s for the simulated separation angles of 55°, 50° and 45°, respectively. The AFD values converge to the AFD of the other separation angles as the shape factor increases. As shown in the angular characteristics results (Fig. 4), when the response function becomes sharper, the drop points of AFD for small separation angles indicate the boundaries at which CSD is just able to separate the two fiber populations. In case of the 40° separation angle, only one FOD peak is obtained. The large difference in AFD for small separation angles (45°-55°) with decreased shape factors can be a confounding factor in inter-subject comparisons of AFD studies, which will be discussed further in Section 4.3. At SNR 10, the AFD differences are more related to noise than to the shape of the response function for smaller separation angles (below 60°). As for the second peak (Fig. 6B), the AFD can change from -30% to 20% when the shape factor was modified from -50% to 50%, respectively. Fig. 6C and Fig. 6D show the percentage difference of the AFD of the first and second fiber population in relation to the scaling factor of the response function. In line with the simulation results for single fiber populations (Fig. 2D), AFD can change up to 80% due to the scaling factor changes for the second peak. For simulated separation angles of approximately 45°, AFD of the first fiber population can be over-estimated up to as much as 150%. For the other simulated separation angles, the AFD of the primary peak can vary from -40% to 70% at SNR = 50 and SNR = 30, irrespective of the simulated separation angles. Notice that the AFD changes are not linearly related with changes in the scaling factor.

**Fig. 6.**
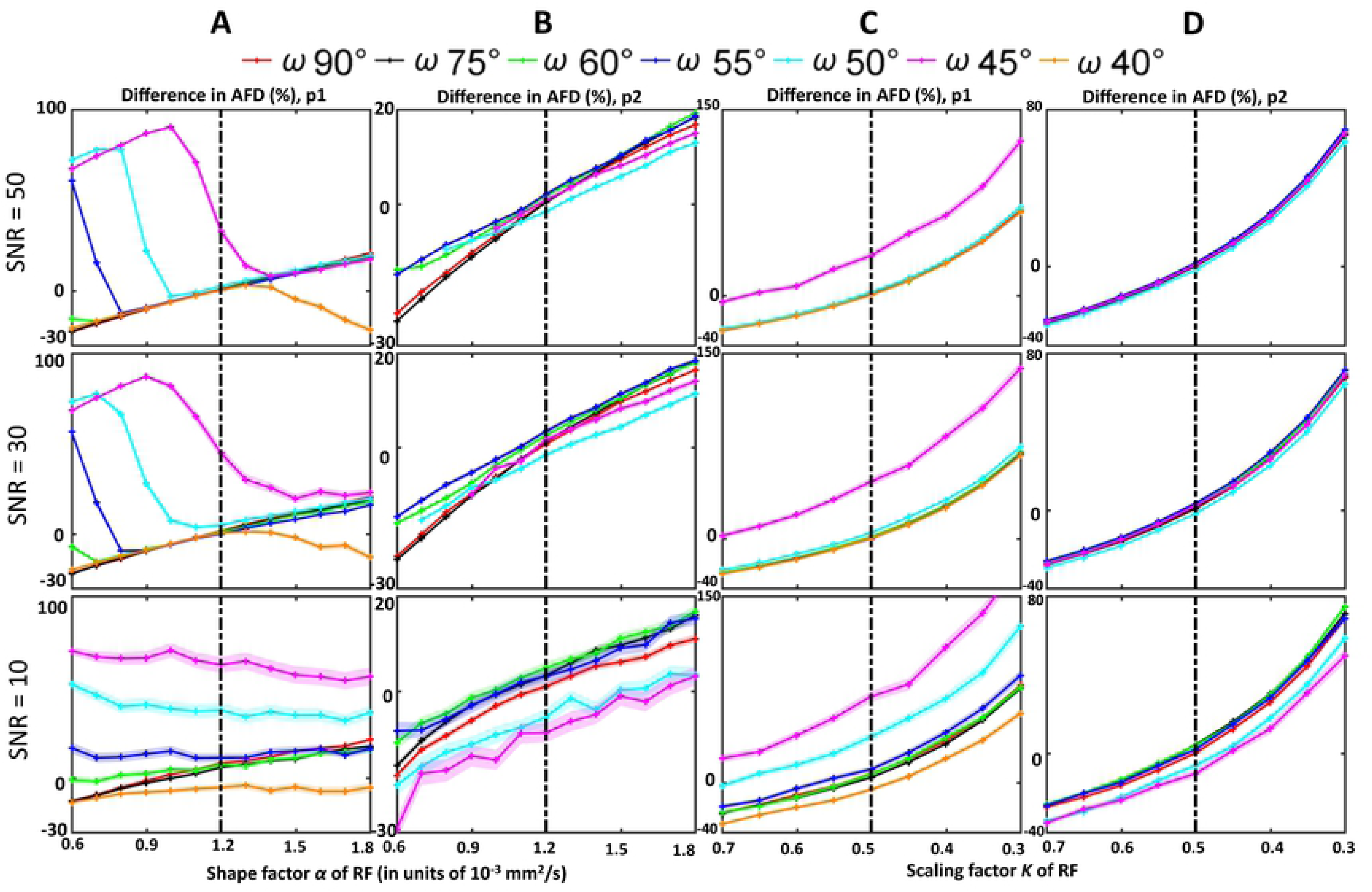
The percentage difference of the estimated AFD of the first peak (p1) and the second peak (p2) in relation to the response function shape factor α (A, B) and scaling factor *K* (C, D) at different SNR levels. The quick drop of the AFD difference while increasing the shape factor indicates when CSD was able to separate the two fiber populations. Dashed vertical lines are defined as in Fig. 3.

### 3.5 In vivo HCP data set

#### 3.5.1 FOD characteristics of white matter

In this section, we present the effect of changing the shape and scaling factors of the response function on FOD characteristics for an axial slice of the HCP data set. The difference in number of FOD peaks per voxel is shown in Fig. 7. Differences are typically seen in areas with partial volume effects and with mostly a peak number difference value of one. When the difference in shape factor, denoted by Δα, increases by 0.1 × 10 ^‒ 3^ mm^2^/s to 0.3 × 10 ^‒ 3^ mm^2^/s, one can see the increase in occurrence of peak number deviations, such as, for instance, in mid-sagittal regions of the corpus callosum. With the increase of difference in scaling factor, denoted by Δ*K*, regions containing CSF showed higher peak number differences than regions with white and gray matter.

**Fig. 7.**
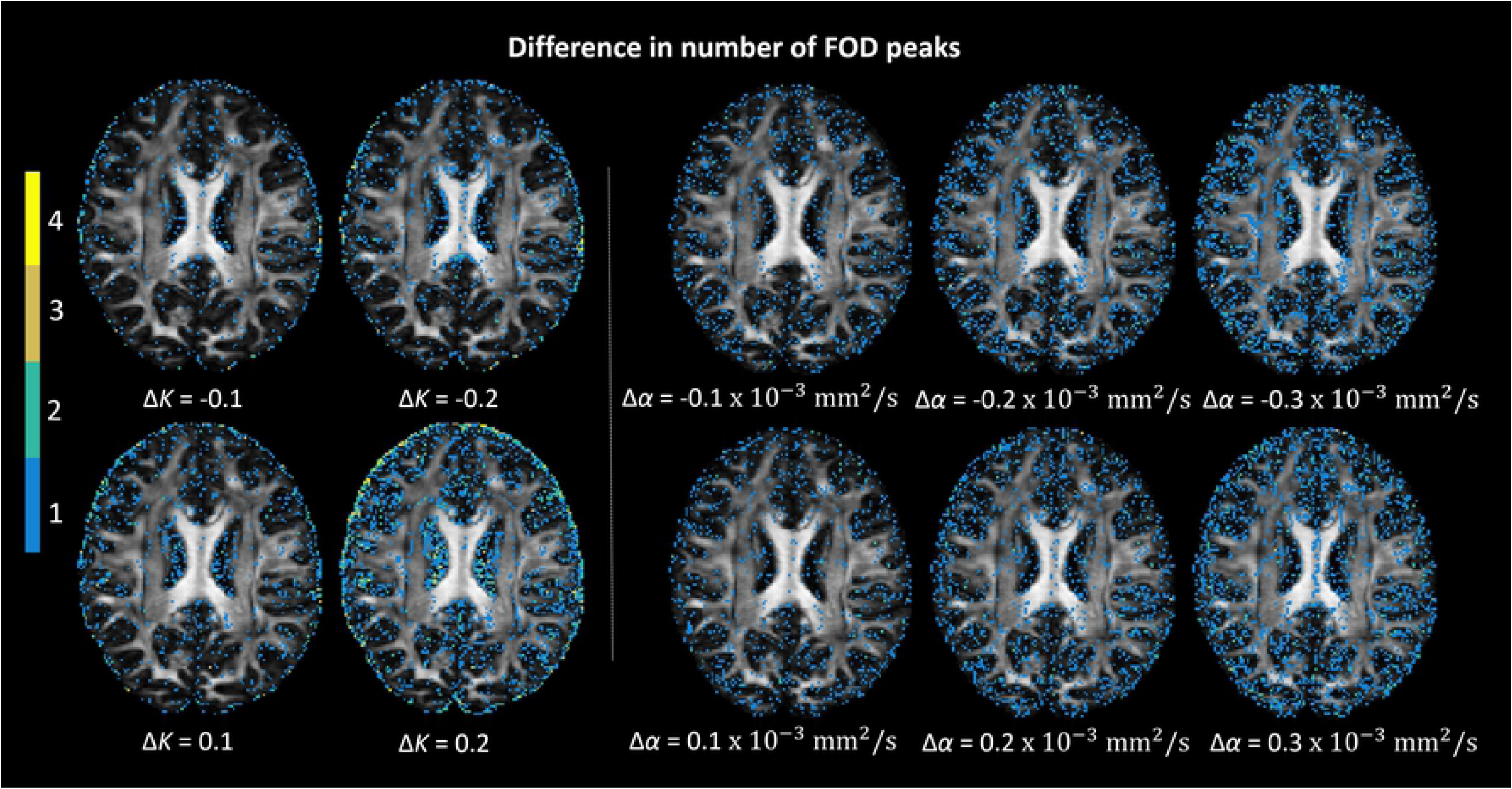
The difference between the number of FOD peaks estimated with the tensor-based response function and the number of FOD peaks computed with the response function modified according to certain changes in scaling (Δ*K*) and shape (Δα) factors. The background is an axial view of the FA map. The peak number difference mostly occurs in grey matter and CSF areas, and crossing fiber regions for white matter, as indicated by the colormap. In regions with single fiber populations (e.g., middle parts of the corpus callosum) spurious peaks are hardly present.

Fig. 8 shows the angular difference between the primary FOD peak, computed with the tensor-fit to the recursive calibrated response function, and the FOD peak obtained with the modified shape and scale factors of the response function. In general, regions containing crossing fibers are affected most when modifying the shape of response functions, with angular deviations of the main FOD peak of up to 3°. Notice that the angular deviation is mostly affected by changing the shape factor, rather than the scaling factor. In addition, while changing Δ*K* did not affect the angular deviation, increasing the magnitude of Δα resulted in larger angular deviations in the same locations.

**Fig. 8.**
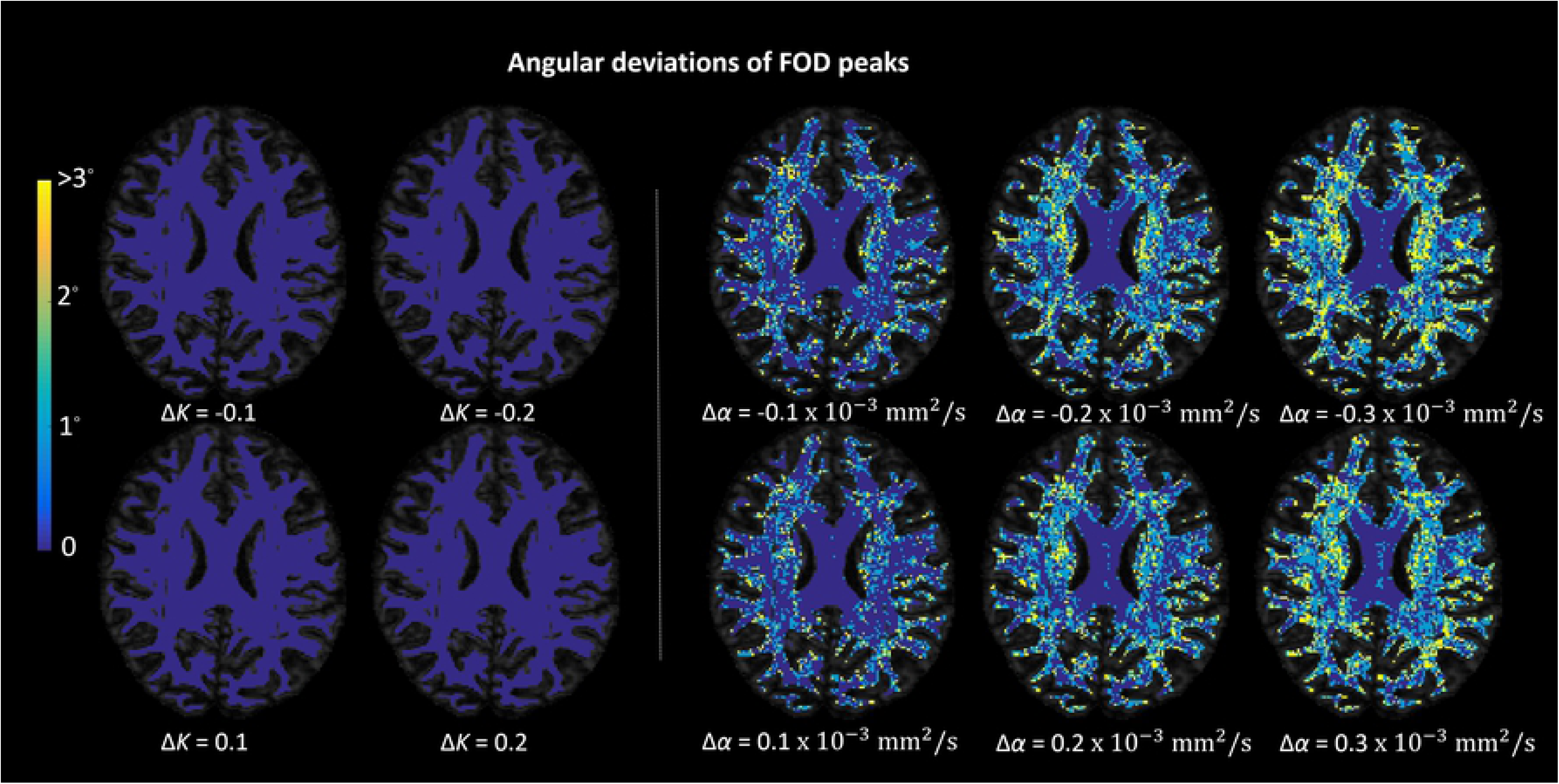
The angular deviations between the FOD peaks estimated with the tensor-fit of the response function and the FOD peaks estimated with the response function modified according to certain changes in scaling (Δ*K*) and shape (Δα) factors. The background is an axial view of the FA map and, for clarity, the angular deviations are shown only in regions where FA > 0.2. Most angular differences are in the range of 0-3°. Similar to the results of spurious peaks shown in Fig. 7, angular deviations are larger in regions with crossing fiber populations than regions with single fiber populations, such as the middle part of the corpus callosum. Notice that the angular deviations are much higher with regard to shape factor changes than scaling factor changes.

Fig. 9 shows the voxel-wise AFD difference for the dominant fiber direction between the FOD estimated using the tensor-fit to the recursive calibrated response function and the FOD obtained with the modified shape and scale factors of the response function for the HCP data set. The AFD shows a very different pattern in relation to the shape factor changes compared to scaling factor changes. The AFD differences are homogenous throughout the brain when the scaling factor varies, while the outliers indicate the voxels where there are potential geometrical differences in the estimated AFD from the reference, such as merging or spurious peaks. The AFD differences are up to 98% when the scaling factor *K* decreased by 0.2. When changing the shape factor with -0.3 × 10 ^‒ 3^ mm^2^/s to 0.3 × 10 ^‒ 3^ mm^2^/s, the highest differences (around 6 to 8%) were observed in areas with a single-fiber population, such as the corpus callosum. Notice that bigger changes of the shape factor *α* makes the AFD difference more heterogeneous across the brain.

**Fig. 9.**
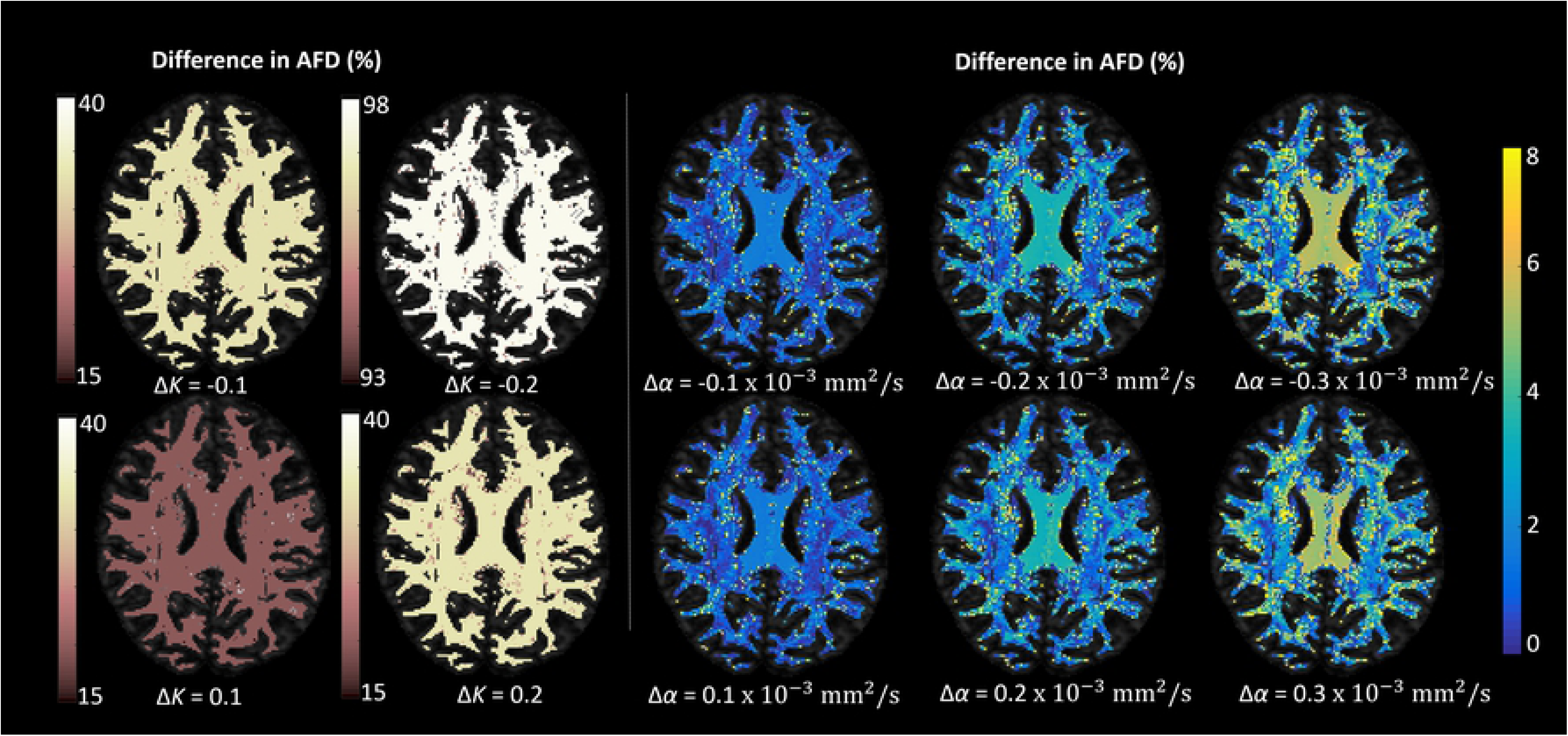
The percentage difference of the apparent fiber density (AFD) between the FOD peaks estimated with the tensor-fit of the response function and the FOD peaks estimated with the response function modified according to certain changes in scaling (Δ*K*) and shape (Δα) factors. The background is an axial view of the FA map and, for clarity, the AFD percentage differences are shown only in regions where FA > 0.2. Notice that the AFD difference stays homogenous with respect to the scaling factor changes, whereas it is heterogeneous when the shape factor changes.

#### 3.5.2 Effect on fiber tractography

Fig. 10 shows the effect of changing the scaling and shape factors of the response function on the reconstruction of the pathways of the tSLF. The reference trajectories (shown in yellow) are computed with the recursive calibration method. While not much differences can be observed for the main part of the reconstructed tracts, changing the response function mainly affected the trajectories where the tSLF enters the frontal and temporal lobes (see enlarged regions in Fig. 10).

**Fig. 10.**
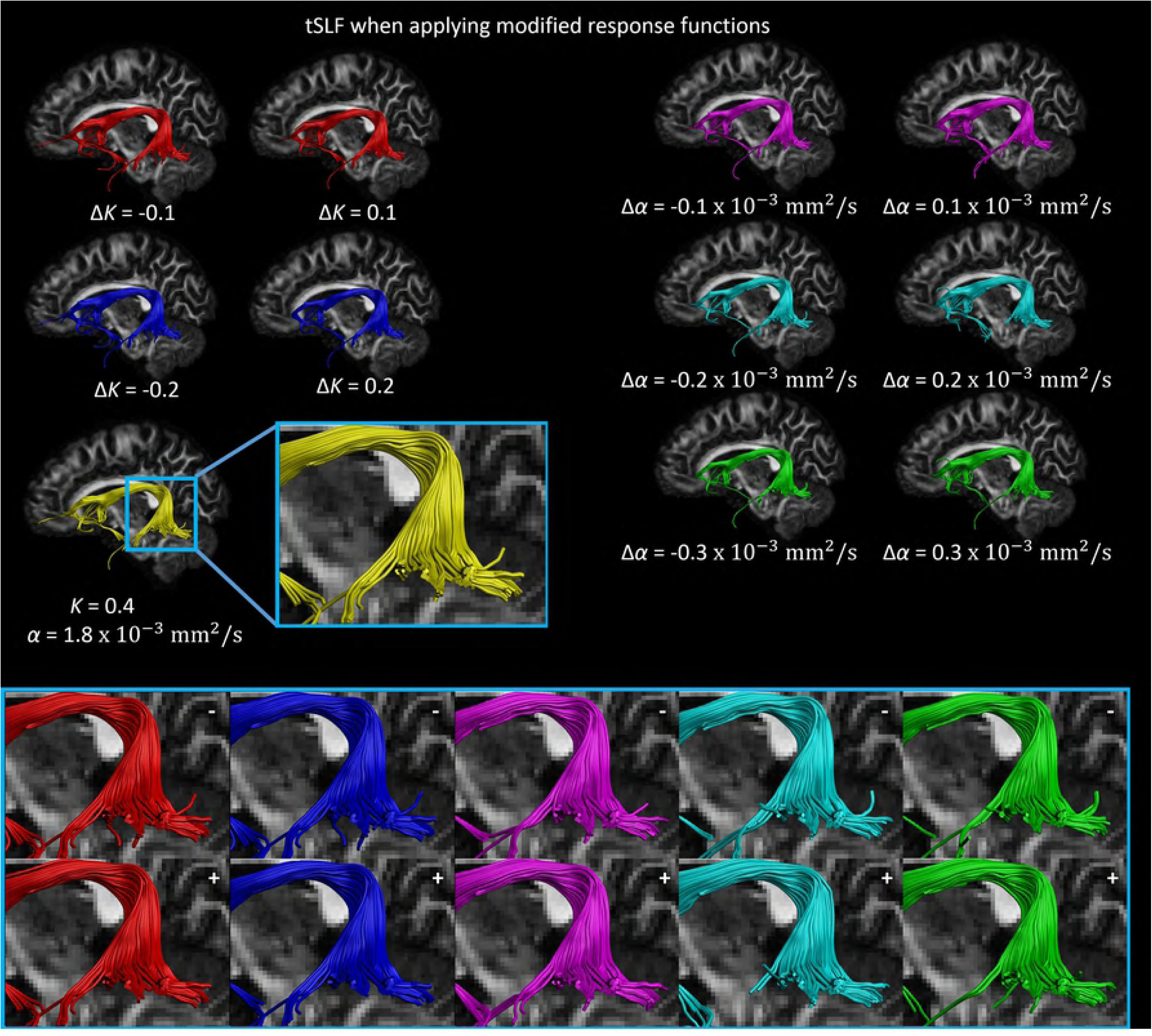
The temporal part of the superior longitudinal fasciculus (tSLF) reconstructed with the FODs estimated using the tensor-fit to the recursively calibrated response function (yellow), and the tSLF from the same ROIs reconstructed with FODs estimated using the modified response functions. The other fiber bundles (shown in red, blue, cyan, magenta, and green) indicate the effect of changing the scaling (Δ*K*) and shape (Δα) factors of the response function on the trajectory of the tSLF. Notice the subtle differences in how the fiber trajectories terminate in the temporal lobe (zoomed areas; the “+” and “-” indicate increase and decrease in the scaling and shape factors, respectively).

Fig. 11 shows the FOD characteristics for the FA-mask, the SFP-mask, and the extracted fiber bundles (gCC, sCC, CST, UF, Cg and tSLF). From all the three FOD characteristics (i.e., spurious peaks, angular deviations, and AFD percentage differences), we can spot a similar trend for all the bundles and the masks with respect to the changes in the shape and scaling factors of the response function. Overall, the UF has the highest average number of spurious peaks. The lowest average angular deviations of the first FOD peak can be seen for the SFP-mask. Furthermore, the alterations of the shape factor of the response function can cause angular deviations up to 6°, while the alterations of the scaling factor hardly cause any angular differences in the masks or the selected fiber bundles (see the enlarged plot). Finally, the differences in AFD are relatively homogenous across the extracted fiber bundles and masks with as a function of changing the shape or the scaling factors.

**Fig. 11.**
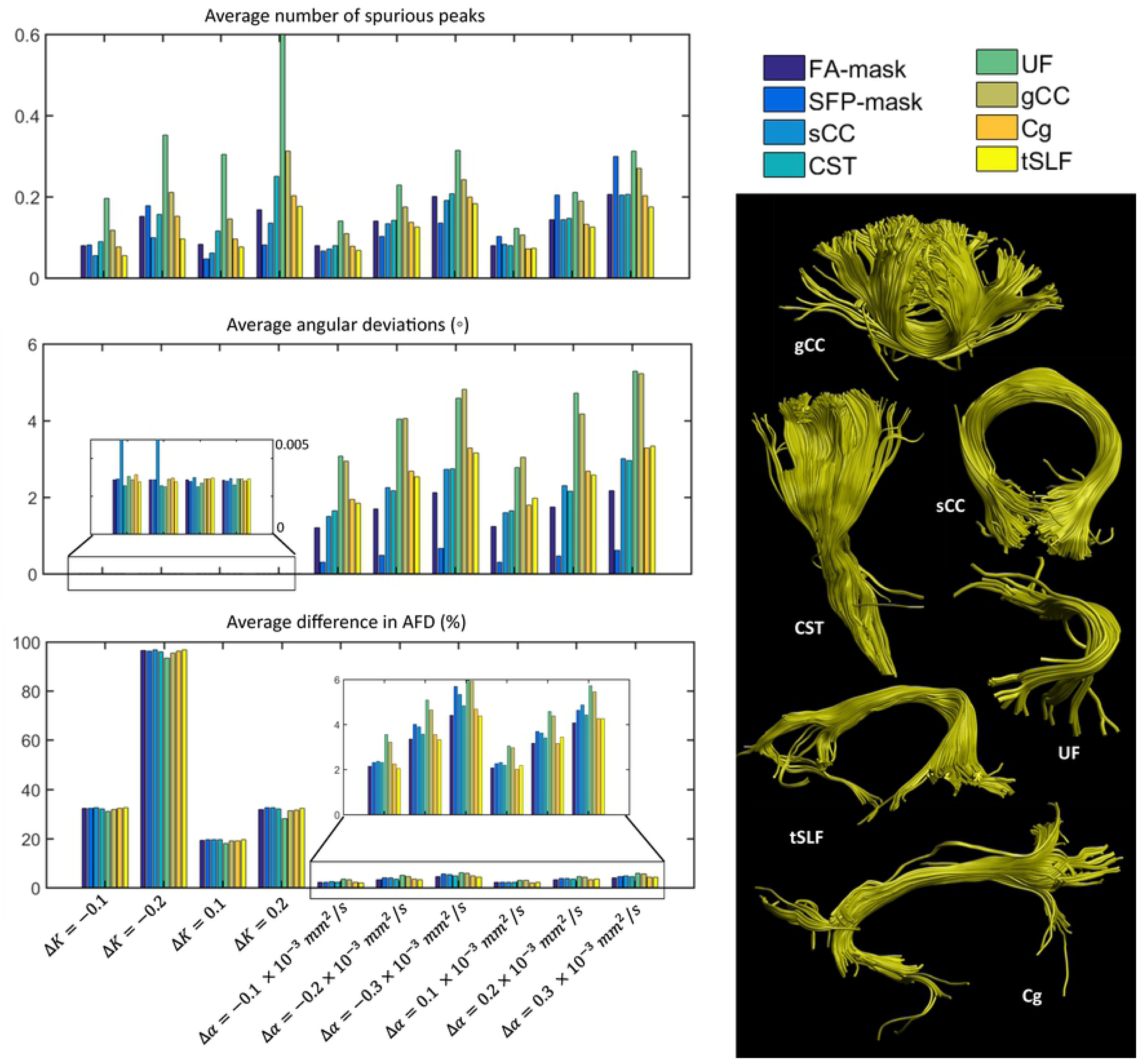
The average number of spurious peaks, the average angular deviations, and the average percentage differences in AFD of the first fiber population for the FA-mask, the SFP-mask, and the selected fiber bundles (shown on the right) when a modified response function was used in comparison to the original tensor-fit to the recursive calibrated response function. The effect of the changes in the scaling (Δ*K*) and shape (Δα) factors of the response function on the selected fiber bundles are reflected in the different color encoding. sCC = splenium of corpus callosum; gCC = genu of corpus callosum; Cg = cingulum; UF = uncinate fasciculus; CST = corticospinal tract; tSLF = temporal part of superior longitudinal fasciculus.

## 4. Discussion

In this work we investigated the effect of changing response function properties on the FOD characteristics using numerical simulations and in vivo HCP data. In particular, we show how miscalibration of the response function, as defined by adjusting the scaling and shape factors, can introduce a bias in the orientation and magnitude of fiber population peaks. Our findings demonstrate that CSD is prone to produce spurious FOD peaks in the presence of miscalibrated response functions, especially in data with insufficient SNR levels. The occurrence of such spurious peaks can also introduce inaccurate fiber pathway reconstructions with fiber tractography. Overall, in agreement with former studies, spurious peaks are introduced due to overestimating the shape factor of the response function, while underestimating the shape factor will result in lower angular resolution of the FOD lobes [10, 15]. Proper tuning of the response function is therefore necessary to achieve an optimal balance between increasing the angular resolution and minimizing the number of spurious peaks, especially for smaller separation angles (i.e., below 60°) and at low SNR levels. Further, AFD estimation can be influenced by the choice of response function, which will be discussed in Section 4.3.

### 4.1 Effect of shape and scaling factors with simulations

At SNR levels of 30 and 50, the FOD characteristics are consistently affected by the choice of the response functions, while at SNR of 10, noise is the dominating factor that affects the FOD properties (Fig. 3). In addition, more spurious peaks are observed at SNR of 10. At relatively high SNR levels, the shape factor of the response function has a greater impact on the results than the scaling factor. In particular, using a sharper response function for separation angles below 50° can potentially increase the angular resolution of CSD and can, therefore, improve the estimation of the number of peaks (Fig. 3). The shape of the response function was reported to vary with axonal injury and brain maturation, whereas the scaling factor was observed to change as result of demyelination, axonal diameters and axonal density changes [10, 35]. This implies that in brain regions affected by disease, applying CSD with a response function determined by healthy white matter data can result in unreliable estimates of FOD characteristics.

### 4.2 Effect of the separation angle between crossing fiber populations

The extent to which the FODs will be affected by the response function depends largely on the separation angle between crossing fiber populations (Fig. 4). More orthogonally crossing fiber orientations are less sensitive to response function changes, as originally suggested in the spherical deconvolution paper [6]. In voxels containing crossing fiber configurations with smaller separation angles (e.g., below 60°), the average angular deviations and their variance increase rapidly with lower shape factors of the response function. By contrast, a higher shape factor of the response function results in a smaller bias in the computation of the FOD peak orientations than the underestimation of the shape factor (Fig. 4 and Fig. 5).

### 4.3 Effect of shape factor on AFD

For fiber populations with separation angles below 55°, CSD fails to estimate the correct number of peaks when response functions with a lower shape factor are employed, leading to artificially higher AFD values (Fig. 6). As FOD peaks merge together when the shape factor is further decreased, the AFD becomes close to the integral of the total FOD amplitudes within the voxel. This is shown in Fig. 6 for simulated separation angles between 45° to 55°. For these relatively small separation angles, the large AFD difference is caused by the limited angular resolution of CSD with the simulated settings. Previous studies [36] reports AFD as a more sensitive diffusion marker in traumatic brain injury than the traditional metrics. However, one should be aware that these changes in AFD in the presence of pathology could result from global response function differences between subjects, rather than local diffusivities alterations.

### 4.4 Effect of FOD angular deviations on fiber tracking

If the angular deviations of the FOD peaks are similar in the neighborhood voxels along the white matter pathways, accumulating effects on reconstructed fibers will be significant. By contrast, the heterogonous angular deviations of the FOD peaks may only change the voxel-wise characteristics like AFD and number of fiber population peaks, the fiber pathways remains if the angular deviations of FOD was not big enough to end in different voxels in the trajectory. Generally, fiber tractography results will not be severely affected in the main part of the fiber bundles, but may show subtle differences at the edges (Fig. 10). In addition, the termination of fiber pathways passing through crossing regions can be affected [12]. With the in vivo HCP data, only minor changes in the tSLF trajectories are detected when using the modified response functions with different shape factors. Nevertheless, an inaccurate response function will influence the FODs and subsequently fiber tractography results.

### 4.5 Limitations and future directions

The reference value of the shape and scaling factor of the simulated diffusion-weighted signals match with the values in the corpus callosum as reported before. However, recent studies [37–40] indicated that the diffusivities of fiber bundles in the brain are not always the same. There is not a full map of diffusivity characteristics of each white matter structure yet. Although our simulation study included the same configurations of crossing fiber bundles in a voxel, in reality, the diffusivities of these crossing fibers may not be identical.

In this study, we showed tractography results of an HCP subject using the tensor-fit to the recursively calibrated response function and modified response functions. In group studies between healthy subjects and patients with neural degradation diseases (e.g., Alzheimer’s disease), it would be useful to compare the alterations of response functions. If there is a group-wise alteration of the shape and the scaling factor of the response functions, we should first exclude the deviations of the diffusivities of the diseased group from the healthy subjects, to ensure that FOD characteristics and fiber tractography changes are not the effects of the response function alteration itself. Furthermore, we can separate the effects of disease on white matter fiber tracking from the effects of response functions used in the FOD estimation.

## 5. Conclusion

This study demonstrates with numerical simulations and in vivo HCP data that decreasing the shape factor of the response function can cause large angular deviations of the FOD peak orientations in crossing fibers. Sharper response functions are responsible for introducing spurious peaks, which can also confound subsequent tractography results. Extremely low shape factors of the response function can cause significant angular deviations and may complicate the interpretation in studies involving pathology. In addition, although individual angular deviations of FOD peak orientations are small for single voxels at most separation angles, the adverse effect can accumulate for brain tractography. Since smaller separation angles are more sensitive to changes of response function shape factors, future work of inter-subject AFD and pathological groups should be aware of this possible confounding factor when investigating brain structures with crossing fiber configurations.

## References

1. Stejskal EO, Tanner JE. Spin diffusion measurements: Spin echoes in the presence of a time-dependent field gradient. J Chem Phys. 1965;42(1):288–92.

2. Le Bihan D, Breton E, Lallemand D, Grenier P, Cabanis E, Laval-Jeantet M. MR imaging of intravoxel incoherent motions: application to diffusion and perfusion in neurologic disorders. Radiology. 1986;161:401–7.

3. Basser PJ, Mattiello J, LeBihan D. MR diffusion tensor spectroscopy and imaging. Biophys J [Internet]. 1994;66(1):259–67. Available from: http://dx.doi.org/10.1016/S0006-3495(94)80775-1

4. Jeurissen B, Leemans A, Tournier JD, Jones DK, Sijbers J. Investigating the prevalence of complex fiber configurations in white matter tissue with diffusion magnetic resonance imaging. Hum Brain Mapp. 2013;34(11):2747–66.

5. Behrens TEJ, Berg HJ, Jbabdi S, Rushworth MFS, Woolrich MW. Probabilistic diffusion tractography with multiple fibre orientations: What can we gain? Neuroimage. 2007;34:144– 55.

6. Tournier JD, Calamante F, Gadian DG, Connelly A. Direct estimation of the fiber orientation density function from diffusion-weighted MRI data using spherical deconvolution. Neuroimage. 2004;23(3):1176–85.

7. Tournier JD, Calamante F, Connelly A. Robust determination of the fibre orientation distribution in diffusion MRI: Non-negativity constrained super-resolved spherical deconvolution. Neuroimage. 2007;35:1459–72.

8. Dell’Acqua F, Rizzo G, Scifo P, Clarke RA, Scotti G, Fazio F. A model-based deconvolution approach to solve fiber crossing in diffusion-weighted MR imaging. IEEE Trans Biomed Eng. 2007;54(3):462–72.

9. Raffelt D, Tournier JD, Rose S, Ridgway GR, Henderson R, Crozier S, et al. Apparent Fibre Density: A novel measure for the analysis of diffusion-weighted magnetic resonance images. Neuroimage [Internet]. 2012;59(4):3976–94. Available from: http://dx.doi.org/10.1016/j.neuroimage.2011.10.045

10. Dell’Acqua F, Simmons A, Williams SCR, Catani M. Can spherical deconvolution provide more information than fiber orientations? Hindrance modulated orientational anisotropy, a true-tract specific index to characterize white matter diffusion. Hum Brain Mapp. 2013;34(10):2464–83.

11. Dhollander T, Raffelt D, Connelly A. Unsupervised 3-tissue response function estimation from single-shell or multi-shell diffusion MR data without a co-registered T1 image. ISMRM Work Break Barriers Diffus MRI [Internet]. 2016;5. Available from: https://www.researchgate.net/publication/307863133_Unsupervised_3-tissue_response_function_estimation_from_single-shell_or_multi-shell_diffusion_MR_data_without_a_co-registered_T1_image

12. Tax CMW, Jeurissen B, Vos SB, Viergever M a., Leemans A. Recursive calibration of the fiber response function for spherical deconvolution of diffusion MRI data. Neuroimage [Internet]. 2014;86:67–80. Available from: http://dx.doi.org/10.1016/j.neuroimage.2013.07.067

13. Jeurissen B, Sijbers J, Tournier J-D. Assessing inter-subject variability of white matter response functions used for constrained spherical deconvolution. In: ISMRM 23th Annual Meeting, Toronto, Ontario, Canada. 2015. p. 2834.

14. Van Essen DC, Ugurbil K, Auerbach E, Barch D, Behrens TEJ, Bucholz R, et al. The Human Connectome Project: A data acquisition perspective. Neuroimage. 2012;62(4):2222–31.

15. Parker GD, Marshall D, Rosin PL, Drage N, Richmond S, Jones DK. A pitfall in the reconstruction of fibre ODFs using spherical deconvolution of diffusion MRI data. Neuroimage [Internet]. 2013;65:433–48. Available from: http://dx.doi.org/10.1016/j.neuroimage.2012.10.022

16. Wheeler-Kingshott CAM, Cercignani M. About “axial” and “radial” diffusivities. Magn Reson Med. 2009;61(5):1255–60.

17. Dell’Acqua F, Scifo P, Rizzo G, Catani M, Simmons A, Scotti G, et al. A modified damped Richardson-Lucy algorithm to reduce isotropic background effects in spherical deconvolution. Neuroimage [Internet]. 2010;49(2):1446–58. Available from: http://dx.doi.org/10.1016/j.neuroimage.2009.09.033

18. Anderson AW. Measurement of fiber orientation distributions using high angular resolution diffusion imaging. Magn Reson Med. 2005;54:1194–206.

19. Jian B, Vemuri BC. A unified computational framework for deconvolution to reconstruct multiple fibers from diffusion weighted MRI. IEEE Trans Med Imaging. 2007;26(11):1464–71.

20. Descoteaux M, Angelino E, Fitzgibbons S, Deriche R. Regularized, fast, and robust analytical Q-ball imaging. Magn Reson Med. 2007;58:497–510.

21. Tuch DS. Q-ball imaging. Magn Reson Med. 2004;52(6):1358–72.

22. Jansons KM, Alexander DC. Persistent angular structure: new insights from diffusion magnetic resonance imaging data. Inverse Probl [Internet]. 2003;19(5):1031. Available from: http://stacks.iop.org/0266-5611/19/i=5/a=303

23. Wedeen VJ, Hagmann P, Tseng WYI, Reese TG, Weisskoff RM. Mapping complex tissue architecture with diffusion spectrum magnetic resonance imaging. Magn Reson Med. 2005;54(6):1377–86.

24. Ramirez-Manzanares A, Rivera M, Vemuri BC, Carney P, Mareei T. Diffusion basis functions decomposition for estimating white matter intravoxel fiber geometry. IEEE Trans Med Imaging. 2007;26(8):1091–102.

25. Tournier J-D, Calamante F, Connelly A. MRtrix: Diffusion tractography in crossing fiber regions. Int J Imaging Syst Technol [Internet]. 2012;22(1):53–66. Available from: http://dx.doi.org/10.1002/ima.22005

26. Garyfallidis E, Brett M, Amirbekian B, Rokem A, Van Der Walt S, Descoteaux M, et al. Dipy, a library for the analysis of diffusion MRI data. Front Neuroinform. 2014;8:8.

27. Leemans A, Jeurissen B, Sijbers J, Jones D. ExploreDTI: a graphical toolbox for processing, analyzing, and visualizing diffusion MR data. Proc 17th Sci Meet Int Soc Magn Reson Med [Internet]. 2009;17(2):3537. Available from: http://www.mendeley.com/research/exploredti-a-graphical-toolbox-for-processing-analyzing-and-visualizing-diffusion-mr-data/%5Cnhttp://www.exploredti.com/ref/ExploreDTI_ISMRM_2009.pdf

28. Jeurissen B, Leemans A, Jones DK, Tournier J-D, Sijbers J. Probabilistic fiber tracking using the residual bootstrap with constrained spherical deconvolution. Hum Brain Mapp. 2011 Mar;32(3):461–79.

29. Jeurissen B, Leemans A, Tournier J-D, Jones DK, Sijbers J. Investigating the prevalence of complex fiber configurations in white matter tissue with diffusion magnetic resonance imaging. Hum Brain Mapp. 2013;34(11):2747–66.

30. Smith RE, Tournier JD, Calamante F, Connelly A. SIFT: Spherical-deconvolution informed filtering of tractograms. Neuroimage [Internet]. 2013;67:298–312. Available from: http://dx.doi.org/10.1016/j.neuroimage.2012.11.049

31. Jones DK, Horsfield MA, Simmons A. Optimal strategies for measuring diffusion in anisotropic systems by magnetic resonance imaging. Magn Reson Med. 1999;42(3):515–25.

32. Jbabdi S, Behrens TEJ, Smith SM. Crossing fibres in tract-based spatial statistics. Neuroimage [Internet]. 2010;49(1):249–56. Available from: http://dx.doi.org/10.1016/j.neuroimage.2009.08.039

33. Tax CMW, Westin CF, Dela Haije T, Fuster A, Viergever MA, Calabrese E, et al. Quantifying the brain’s sheet structure with normalized convolution. Med Image Anal [Internet]. 2017;39:162–77. Available from: http://dx.doi.org/10.1016/j.media.2017.03.007

34. Wakana S, Caprihan A, Panzenboeck MM, Fallon JH, Perry M, Gollub RL, et al. Reproducibility of quantitative tractography methods applied to cerebral white matter. Neuroimage [Internet]. 2007;36(3):630–44. Available from: http://dx.doi.org/10.1016/j.neuroimage.2007.02.049

35. Feldman HM, Yeatman JD, Lee ES, Barde LHF, Gaman-Bean S. Diffusion tensor imaging: a review for pediatric researchers and clinicians. J Dev Behav Pediatr JDBP. 2010;31(4):346.

36. Wright DK, Johnston LA, Kershaw J, Ordidge R, O’Brien TJ, Shultz SR. Changes in apparent fiber density and track-weighted imaging metrics in white matter following experimental traumatic brain injury. J Neurotrauma. 2017;34(13):2109–18.

37. Tax CMW, Novikov DS, Garyfallidis E, Viergever MA, Descoteaux M, Leemans A. Localizing and Characterizing Single Fiber Populations Throughout the Brain. Proc 23rd Annu Meet ISMRM, Toronto, Canada [Internet]. 2015;59(6):473. Available from: http://scil.dinf.usherbrooke.ca/wp-content/papers/tax-etal-ismrm15a.pdfVN-readcube.com

38. Novikov DS, Jespersen SN, Kiselev VG, Fieremans E. Quantifying brain microstructure with diffusion MRI: Theory and parameter estimation. ArxivOrg [Internet]. 2016;1–38. Available from: http://arxiv.org/abs/1612.02059

39. Jespersen SN, Kroenke CD, Østergaard L, Ackerman JJH, Yablonskiy DA. Modeling dendrite density from magnetic resonance diffusion measurements. Neuroimage. 2007;34(4):1473–86.

40. Kroenke CD, Ackerman JJH, Yablonskiy DA. On the nature of the NAA diffusion attenuated MR signal in the central nervous system. Magn Reson Med. 2004;52(5):1052–9.

